# APpar: automated action potential parameter analysis software for reproducible electrophysiological measurements in neurons

**DOI:** 10.64898/2026.06.10.731446

**Authors:** Dmytro V. Vasylyev, Stephen G. Waxman

## Abstract

Quantitative analysis of action potential (AP) waveforms is central to studies of neuronal excitability, ion channel function, disease mechanisms, and pharmacological modulation. However, AP analysis is still often performed using partially manual workflows, laboratory-specific spreadsheets, or proprietary software environments that can limit reproducibility, transparency, and throughput. Here we present APpar, a freely available, open-source software tool for extracting AP parameters, developed for use with the OriginLab software package Origin/OriginPro.

APpar detects APs from membrane voltage recordings using a user-defined derivative criterion and calculates a comprehensive set of excitability parameters, including resting membrane potential, AP threshold, dV/dt at threshold, overshoot, undershoot, AP amplitude, AP half-amplitude, rise time, decay time, AP duration, AP half-width, AP width at 0 mV, AP area above voltage threshold, dV/dt_MAX_, dV/dt_MIN_, interspike interval for the respective AP. Because AP threshold is a particularly sensitive and method-dependent measurement, APpar includes a TRUE-threshold validation algorithm. After the initial forward dV/dt threshold crossing is identified, the software finds AP overshoot, searches backward to the closest preceding local dV/dt maximum, then searches backward to the user-defined dV/dt crossing and recalculates AP parameters from this validated threshold point.

We validated APpar using APs from dorsal root ganglion neurons current-clamp recordings, including copied identical APs, current-evoked repetitive firing, and long-duration spontaneous firing. The software produced stable measurements from identical copied APs and extracted dynamic changes in AP parameters across repetitive and spontaneous firing sequences. APpar provides a transparent, customizable, and Origin-compatible framework for reproducible AP analysis in neuronal electrophysiology.

**Significance statement:** Action potential waveform analysis is essential for interpreting neuronal excitability, but many AP measurements remain vulnerable to user-dependent threshold placement, manual cursor selection, and inconsistent parameter definitions. APpar, a freely available, open-source software tool, addresses this problem by automating AP detection and parameter extraction within the OriginLab environment widely available to electrophysiology laboratories. The software formalizes definitions of AP threshold, amplitude, duration, half-width, afterhyperpolarization, derivative-based parameters, and firing metrics, and introduces a TRUE-threshold validation algorithm that recalculates AP parameters from a derivative-validated threshold point. This workflow reduces operator-dependent variability while preserving user control over physiologically meaningful detection criteria.

**Highlights:** Automated action potential waveform analysis within OriginLab Origin environments AP threshold validation improves reproducibility of derivative-based threshold detection Extracts action potential kinetics, amplitudes, widths, and dV/dt measurements Open-source workflow supports reproducible neuronal electrophysiology data analysis Validated using repetitive and spontaneous firing in DRG neurons

## Introduction

Action potentials (AP) are the fundamental electrical signals by which neurons encode excitability, transmit information (Adrian and Forbes, 1922), and report the functional state of voltage-gated ion channels (Hodgkin and Huxley, 1952; Kocsis and Waxman, 1983; Renganathan et al., 2001; Blair and Bean, 2002; Bean, 2007; Kole and Stuart, 2012). An example of the complexity of AP waveform analysis is provided by small-diameter dorsal root ganglion (DRG) neurons, that include nociceptors; these cells express multiple sodium, potassium, calcium, and hyperpolarization-activated channels that shape AP initiation, amplitude, duration, afterhyperpolarization, and repetitive firing (Renganathan et al., 2001; Blair and Bean, 2002; Vasylyev et al., 2023) and that have been assessed, not just in vivo and at the channel level, but also in terms of AP generation, as therapeutic targets (Waxman and Zamponi, 2014; Waxman and Vasylyev, 2025). AP parameters are widely used to characterize nociceptor excitability, inherited and acquired pain mechanisms, pharmacological modulation, and the functional contribution of specific ion channels (Cummins et al., 2004; Waxman, 2007; Vasylyev et al., 2014; Dib-Hajj and Waxman, 2019; Jones et al., 2023; Yang and Prescott, 2023; Vaelli et al., 2024; Vasylyev et al., 2024; Alsaloum et al., 2025; Osteen et al., 2025; Stewart et al., 2025).

Despite the central importance of AP waveform analysis, many laboratories still rely on partially manual workflows. A typical analysis may involve visual inspection, manual placement of threshold cursors, separate calculation of AP width or amplitude, transfer of values into spreadsheets, and repeated post-processing steps. These approaches are slow and can introduce operator-dependent variability. Even when commercial software automates portions of analysis, the underlying algorithms may be opaque or difficult to modify, and customization for laboratory-specific questions may be limited.

The problem is particularly acute for AP threshold. AP threshold is not a directly observed physical boundary; it is an operational definition of the voltage at which the regenerative AP upstroke begins according to a chosen criterion (Naundorf et al., 2006; Platkiewicz and Brette, 2010; Brette, 2013). Different studies define threshold by a fixed derivative criterion, a relative derivative change, an inflection point, a second derivative criterion, or visual inspection. This means that values for threshold voltage, AP amplitude, rise time, AP duration, and AP area can all shift depending on how threshold is defined. (Patrick Harty and Waxman, 2007), for example, defined AP onset in small DRG neurons as the point at which AP dV/dt exceeded 5 mV/ms, choosing this value to be approximately twice the maximum baseline noise of the derivative trace. They defined AP amplitude as the difference between AP peak and AP onset. This definition of AP amplitude explicitly links AP amplitude to a threshold/onset rather than to resting membrane potential (RMP). Because alternative definitions are also commonly used in the literature, including the difference between overshoot and RMP and between overshoot and undershoot, APpar additionally reports overshoot, threshold voltage, RMP, and undershoot values for every analyzed AP. This allows users to calculate AP amplitude and related parameters according to their preferred convention.

Recent studies of DRG neuron excitability further emphasize the importance of precise AP quantification. Dynamic clamp studies have explored how different voltage-gated ion channels shape excitability in small DRG neurons, including the stabilizing effects of HCN channels in the context of Nav1.7-dependent hyperexcitability (Vasylyev et al., 2023), the functional interaction between Nav1.8 and Nav1.7 near AP threshold (Vasylyev et al., 2024), and intrinsic limits on the functional effect of Nav1.8 abrogation (Vasylyev et al., 2025). These and other studies (Xie et al., 2024; Jo et al., 2025; Fujita et al., 2026) require reliable measurements of AP threshold, firing probability, AP amplitude, repetitive firing, and subthreshold-to-threshold excitability. The need for standardized, transparent, and reproducible AP analysis tools is therefore not merely technical; it directly affects interpretation of ion channel function in nociceptors.

Several open-source and semi-open electrophysiology tools have addressed related needs. SimplyFire provides an open-source Python-based platform for synaptic event analysis (Mori et al., 2024). ActionPytential and BAPTA provide cardiac AP analysis workflows (Arpadffy-Lovas and Nagy, 2023; Leonov et al., 2023). APsim was developed within the Origin 8.5 LabTalk framework and computes ionic currents based on ion channel kinetic models. It can be used for both action potential-clamp (or voltage-clamp) and current-clamp simulations (Vasylyev et al., 2014; Vasylyev et al., 2025; Vasylyev et al., 2026). These tools demonstrate the value of transparent and user-friendly electrophysiological software, but neuronal AP analysis in Origin remains a common practical workflow that benefits from a dedicated, automated, customizable tool.

Here we present APpar, a software platform for automated AP parameter analysis in OriginLab Origin/OriginPro. APpar was developed first in LabTalk and then converted into an Origin-Python implementation. The software reads time and voltage columns from an Origin worksheet, calculates raw and smoothed dV/dt, detects APs, validates threshold using a backward-search TRUE-threshold algorithm, applies quality filters, and writes structured results to a new workbook. We describe the APpar workflow, define each extracted AP parameter, and validate the software using representative DRG neuron recordings.

## Materials and Methods

### Software environment and implementation

APpar was implemented in two complementary forms: a native LabTalk program and an Origin-Python program launched through a LabTalk wrapper. The LabTalk implementation preserves compatibility with standard Origin workflows. The Python implementation uses numpy and pandas for numerical operations and table handling, and the Origin/OriginPro Python interface for worksheet input and output. The Python version was designed to reproduce the analytical logic of the LabTalk version while improving execution speed, especially for multiple traces or long recordings.

APpar assumes that the active Origin worksheet contains time in column A(x) and one or more membrane voltage traces in subsequent columns. The user runs APpar from the LabTalk command window after activating the source workbook. The Python implementation reads worksheet columns by position rather than long name, which avoids errors when multiple voltage columns share identical long names. The program writes raw and smoothed dV/dt columns back to the source worksheet and creates a new results workbook with one results sheet per analyzed trace. APpar assumes that input and output time values are in seconds and membrane voltage values are in Volts. A template OriginPro file, APpar.opju, presented in Figure 1, is included for reference (see Supplemental Data).

**Figure 1.**
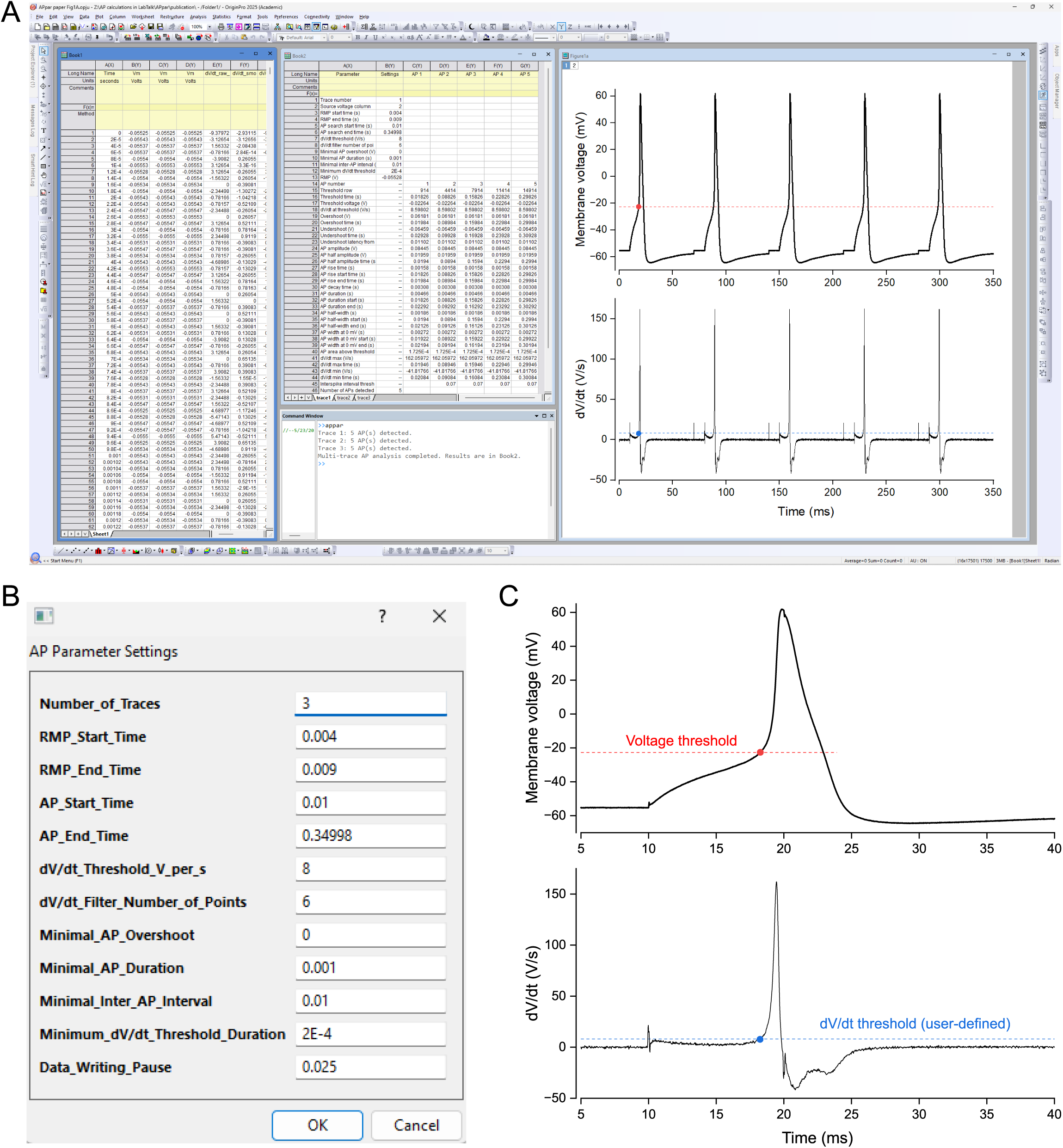
APpar software overview. Representative Origin/OriginPro workflow for APpar. The active worksheet contains time and voltage recordings (A), while the settings dialog window (B) defines analysis parameters including number of traces, RMP interval, AP search interval, dV/dt threshold, derivative smoothing, overshoot filter, AP duration filter, inter-AP interval, and minimum dV/dt threshold duration. Output includes derivative traces and structured AP parameter tables (A, C).

## Installation and execution

APpar was developed and tested under Windows 10 and Windows 11 using Origin 2022b and OriginPro 2025. Two implementations are provided: the Origin-Python implementation, launched by the LabTalk wrapper APpar.ogs, and the LabTalk-only implementation, launched by APparLT.ogs.

For the Origin-Python implementation, the required Python packages are installed inside the Origin/OriginPro Python environment. First, the user opens Origin/OriginPro and selects Connectivity - Python Packages, then installs numpy, scipy, and pandas. Second, the user opens Connectivity - Python Console to verify the active Python environment and install the same packages into that environment. The following commands are entered sequentially, pressing Enter after each line:

import sys

print(sys.executable)

import subprocess

subprocess.check_call([sys.executable, “-m”, “pip”, “install”, “numpy”, “scipy”, “pandas”])

After package installation, Origin/OriginPro should be restarted.

The APpar.ogs file is a LabTalk wrapper that runs APpar.py from Origin/OriginPro while preserving the same settings-dialog style used by the LabTalk implementation. Before running APpar, the user opens APpar.ogs and edits the Python file path so that it matches the local Windows user account. For example, the default line

string pyFile$ = “C:\\Users\\dv93\\Documents\\OriginLab\\User Files\\APpar.py”;

is changed by replacing dv93 with the local Windows user name. Both APpar.ogs and APpar.py are then placed in the OriginLab User Files directory:

C:\Users\USERNAME\Documents\OriginLab\User Files

where USERNAME is the local Windows user name. Origin/OriginPro should be restarted after the files are copied.

For the LabTalk-only implementation, the APparLT.ogs file is copied into the same OriginLab User Files directory:

C:\Users\USERNAME\Documents\OriginLab\User Files

where USERNAME is again replaced with the local Windows user name.

Origin/OriginPro should be restarted after copying APparLT.ogs.

To run either implementation, the user first activates the workbook/worksheet containing the AP trace or traces by clicking on it. The LabTalk Command Window is then opened from Window - Command Window or by pressing Alt+3. Typing APpar and pressing Enter launches the Origin-Python implementation. Typing APparLT and pressing Enter launches the LabTalk-only implementation.

### User-defined settings

APpar uses a compact set of user-defined analysis settings. These include the number of traces, RMP start time, RMP end time, AP search start time, AP search end time, dV/dt threshold, dV/dt smoothing window, minimal AP overshoot, minimal AP duration, minimal inter-AP interval, minimum duration above the dV/dt threshold, and data-writing pause. Default settings in the Python implementation include an RMP window from 0.004 to 0.009 s, AP search start time of 0.01 s, dV/dt threshold of 8 V/s, derivative smoothing over 6 points, minimal overshoot of 0 mV, minimal AP duration of 0.001 s, minimal inter-AP interval of 0.01 s, and minimum dV/dt threshold duration of 0.0002 s. The default settings were optimized for APs recorded from small DRG neurons at room temperature (23 °C); however, the user-friendly parameter settings interface implies broad utility of the program across a wide range of neuronal types and, perhaps, non-neuronal cells.

### AP parameter calculations

The membrane voltage derivative is calculated from the time and voltage vectors. For internal points, APpar uses a symmetric central-difference approximation:

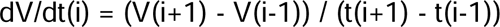

AP half-amplitude time is the time at which the rising phase first reaches this voltage. This value is used to derive AP half-width.

AP rise time is the interval between threshold time and overshoot time. AP rise start time is the threshold time, and AP rise end time is the overshoot time. Rise time is influenced largely by sodium channel availability, activation kinetics, membrane resistance, and recording bandwidth.

AP decay time is the interval between overshoot time and the time at which the membrane voltage returns to threshold voltage during repolarization. This measurement captures the duration of the post-peak decay back to the threshold reference voltage.

AP duration is defined as the interval between validated threshold time and the repolarization crossing of threshold voltage. AP duration start is the validated threshold time, and AP duration end is the time at which the voltage returns to threshold after overshoot. This measurement is distinct from AP half-width because it uses threshold voltage as the reference level.

AP half-width is the time interval between the rising-phase and falling-phase crossings of the AP half-amplitude voltage. AP half-width start is the first time the waveform reaches half amplitude during the rising phase, and AP half-width end is the time the waveform returns to half amplitude during repolarization. Half-width is a widely used measure of spike broadening and is sensitive to sodium current availability, potassium conductance, calcium-activated potassium currents, and cumulative inactivation during repetitive firing.

AP width at 0 mV is the interval between the first upward crossing of 0 mV and the post-overshoot downward crossing of 0 mV. AP width at 0 mV start and end are the corresponding crossing times. This measurement is useful for APs that overshoot 0 mV and provides a standardized suprathreshold width independent of threshold voltage.

AP area above threshold is calculated by numerical integration of the voltage above threshold from threshold time to the return-to-threshold time. This area is reported as ∫(Vm - Vthreshold)dt. It summarizes spike size and duration in a single value and may be sensitive to cumulative waveform changes that are not fully captured by peak or width alone.

dV/dt_MAX_ is the maximum smoothed derivative during the AP, and dV/dt_MAX_ time is the time at which it occurs. This parameter reflects the fastest depolarizing phase of the AP and is commonly used as a proxy for available inward current and AP upstroke speed.

dV/dt_MIN_ is the minimum smoothed derivative during the AP, and dV/dt_MIN_ time is the corresponding time. This parameter reflects the fastest repolarizing phase and is influenced by voltage-gated potassium currents, sodium channel inactivation, and other repolarizing mechanisms.

Number of APs detected is the count of accepted APs in each trace. This is a trace-level summary value and is useful for firing probability and repetitive firing analysis.

Forward threshold row and forward threshold time report the initial derivative crossing detected during the forward scan. TRUE threshold row and TRUE threshold time report the validated threshold after the backward search. Threshold source identifies whether the TRUE threshold was used. Forward versus TRUE row difference reports the absolute sample difference between the initial and validated threshold positions.

### Quality filters

APpar applies quality filters to reduce false-positive detections. Candidate APs are rejected if the overshoot is less than the user-defined minimum, if AP duration is shorter than the user-defined minimum, or if the derivative remains above the threshold criterion for less than the minimum dV/dt threshold duration. The minimal inter-AP interval prevents closely spaced derivative fluctuations from being counted as separate APs.

### Electrophysiology

Whole-cell current-clamp recordings were obtained from rodent small-diameter dorsal root ganglion (DRG) neurons grown in primary culture using methods described previously (Vasylyev et al., 2025). In brief, small DRG neurons (soma diameter 23-30 μm) were recorded in whole-cell current-clamp configuration using a HEKA EPC10 USB amplifier and PatchMaster software (Heka Elektronik, Germany). Voltage and current traces were filtered at 3 kHz and digitized at 50 kHz. Patch pipettes were pulled from borosilicate glass capillaries (World Precision Instruments, Sarasota, FL; PG52165-4) and had resistance of 1.5-3 MΩ when filled with intracellular solution containing (in mM): 150 KCl, 0.5 EGTA, 5 HEPES, 3 Mg-ATP, and 5.6 glucose (pH 7.3 with KOH; 286-288 mOsm). The extracellular solution was HBSS containing (in mM): 138 NaCl, 5.3 KCl, 1.3 CaCl_2_, 0.5 MgCl_2_, 0.4 MgSO_4_, 0.4 KH_2_PO_4_, 0.3 Na_2_HPO_4_, 4.2 NaHCO_3_, and 5.6 glucose (284-286 mOsm). Recordings were performed at room temperature (21-23 °C), and neurons were recorded at native resting membrane potential without steady-state current injection. Membrane voltage recordings were exported into Origin/OriginPro worksheets and analyzed using APpar.

### Validation datasets and figure set

This paper contains seven figures. Figure 1 shows the APpar software overview and parameter settings in Origin/OriginPro. Figure 2 is a newly generated operational diagram and algorithmic workflow. Figure 3 defines AP waveform parameters. Figure 4 illustrates AP threshold determination. Figure 5 tests calculation stability using identical copied APs. Figure 6 shows representative current-evoked AP trains in a small DRG neuron. Figure 7 shows representative spontaneous AP recordings from two small DRG neurons, with parameters calculated across 60 s recordings.

**Figure 2.**
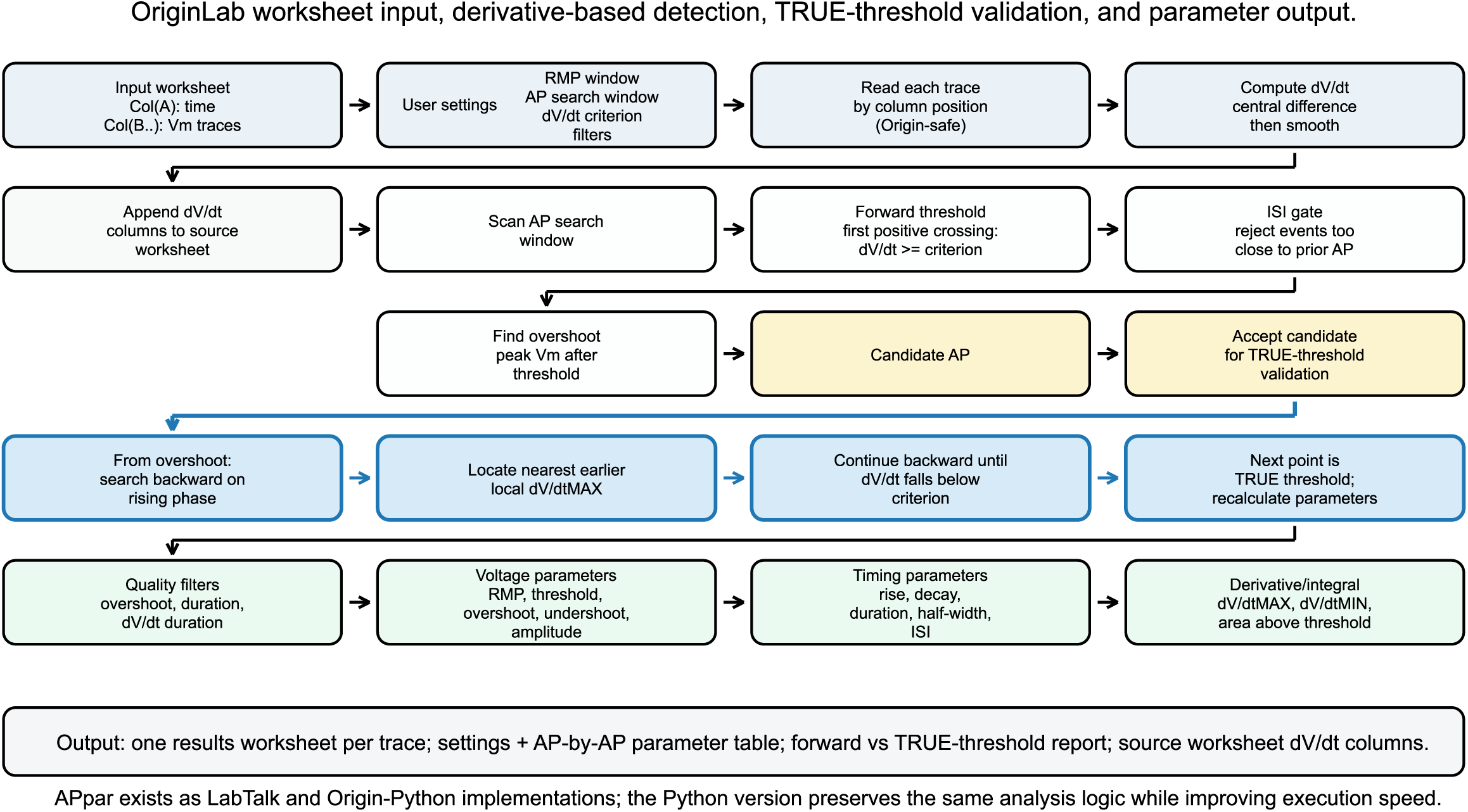
APpar operational workflow and AP detection algorithm. Operational diagram of APpar based on the LabTalk and Origin-Python code structure. The workflow includes worksheet import, dV/dt calculation and smoothing, forward threshold detection, inter-AP interval filtering, overshoot detection, TRUE-threshold validation, AP parameter extraction, quality filtering, and output generation. Blue arrows indicate iterative threshold-validation steps.

**Figure 3.**
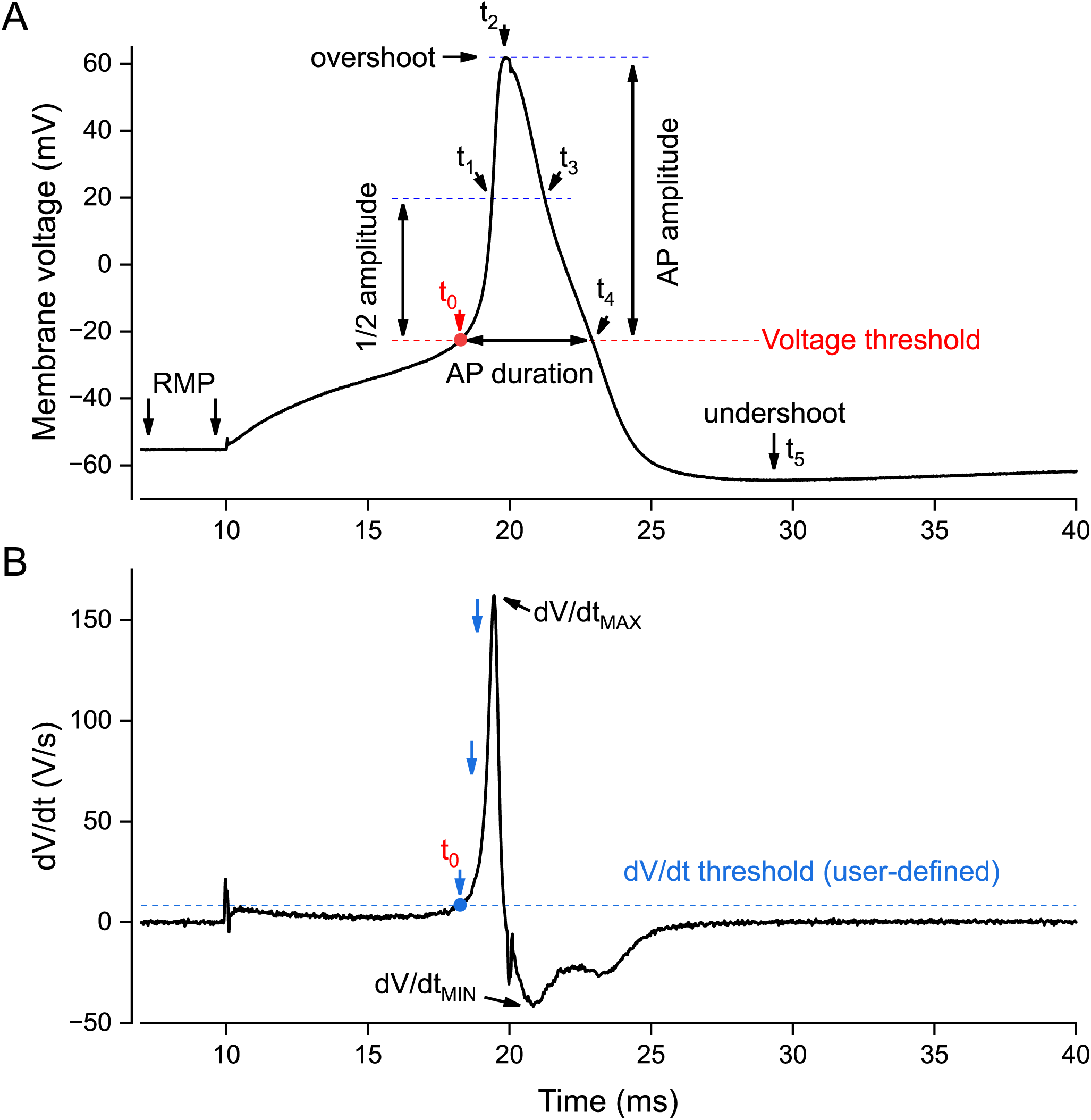
Definition of AP parameters and TRUE-threshold validation. Representative AP waveform (A) and derivative (B) trace showing the parameters calculated by APpar, including threshold, overshoot, undershoot, AP amplitude, AP duration, AP half-width, dV/dt_MAX_, and dV/dt_MIN_. Time points t_0_-t_5_ define key stages of the AP waveform. t_0_ indicates validated AP threshold, t_1_ rising-phase half-amplitude crossing, t_2_ AP overshoot, t_3_ falling-phase half-amplitude crossing, t_4_ return to threshold voltage, and t_5_ AP undershoot. The lower panel (B) illustrates TRUE-threshold validation. After initial forward dV/dt threshold detection, APpar searches backward from AP overshoot to the nearest earlier local dV/dt maximum and then to the user-defined derivative threshold crossing. Blue arrows indicate the evolution of the algorithmic search process.

**Figure 4.**
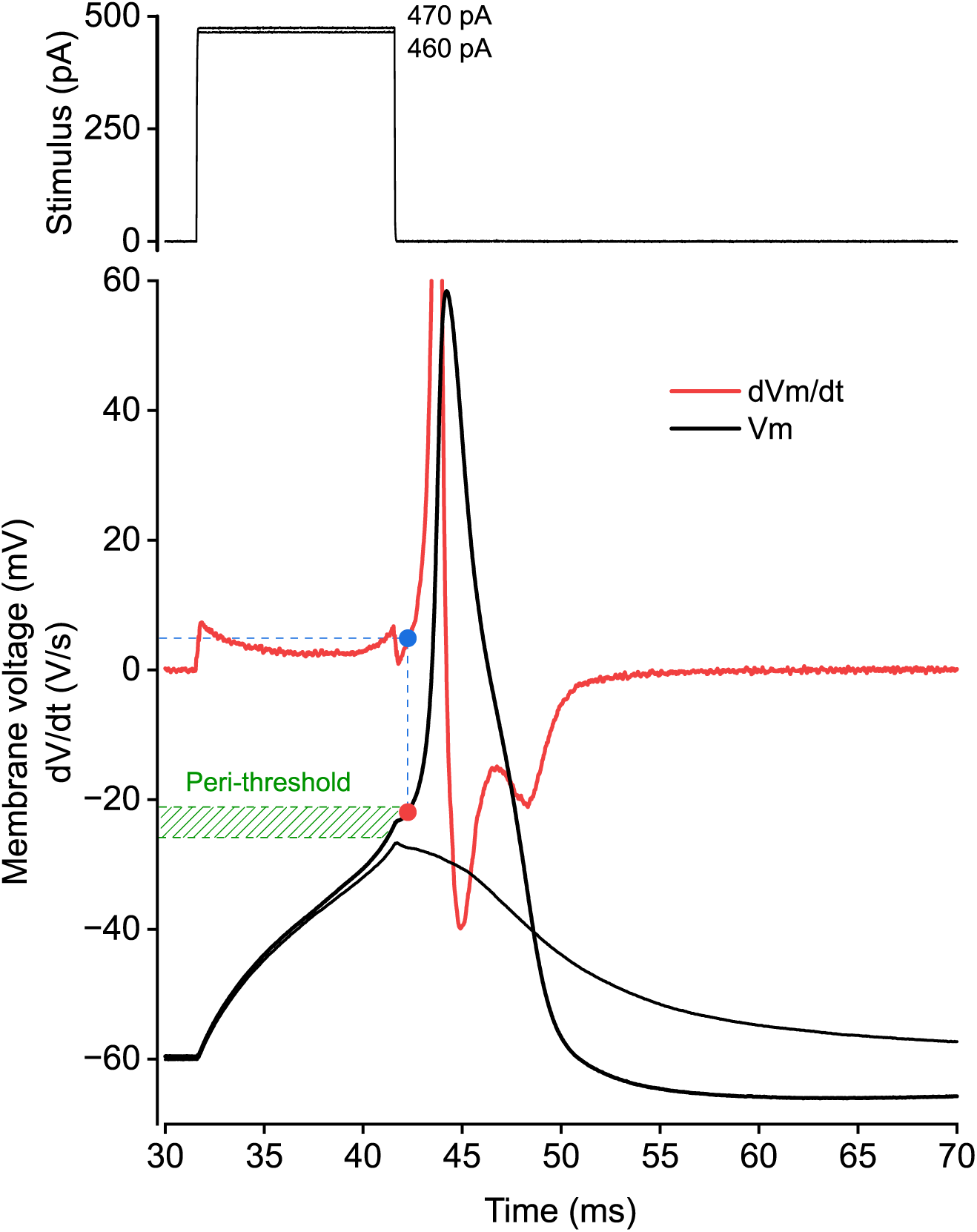
Rheobase, peri-threshold region, and AP threshold voltage. Representative recordings illustrating rheobase determination and peri-threshold membrane behavior in a small DRG neuron. Current injection of 460 pA (upper panel) did not evoke an action potential, whereas 470 pA evoked an AP (lower panel), defining rheobase under these recording conditions. The lower panel shows the evoked AP overlayed with its derivative and highlights the peri-threshold region (green) associated with AP initiation and threshold voltage determination.

**Figure 5.**
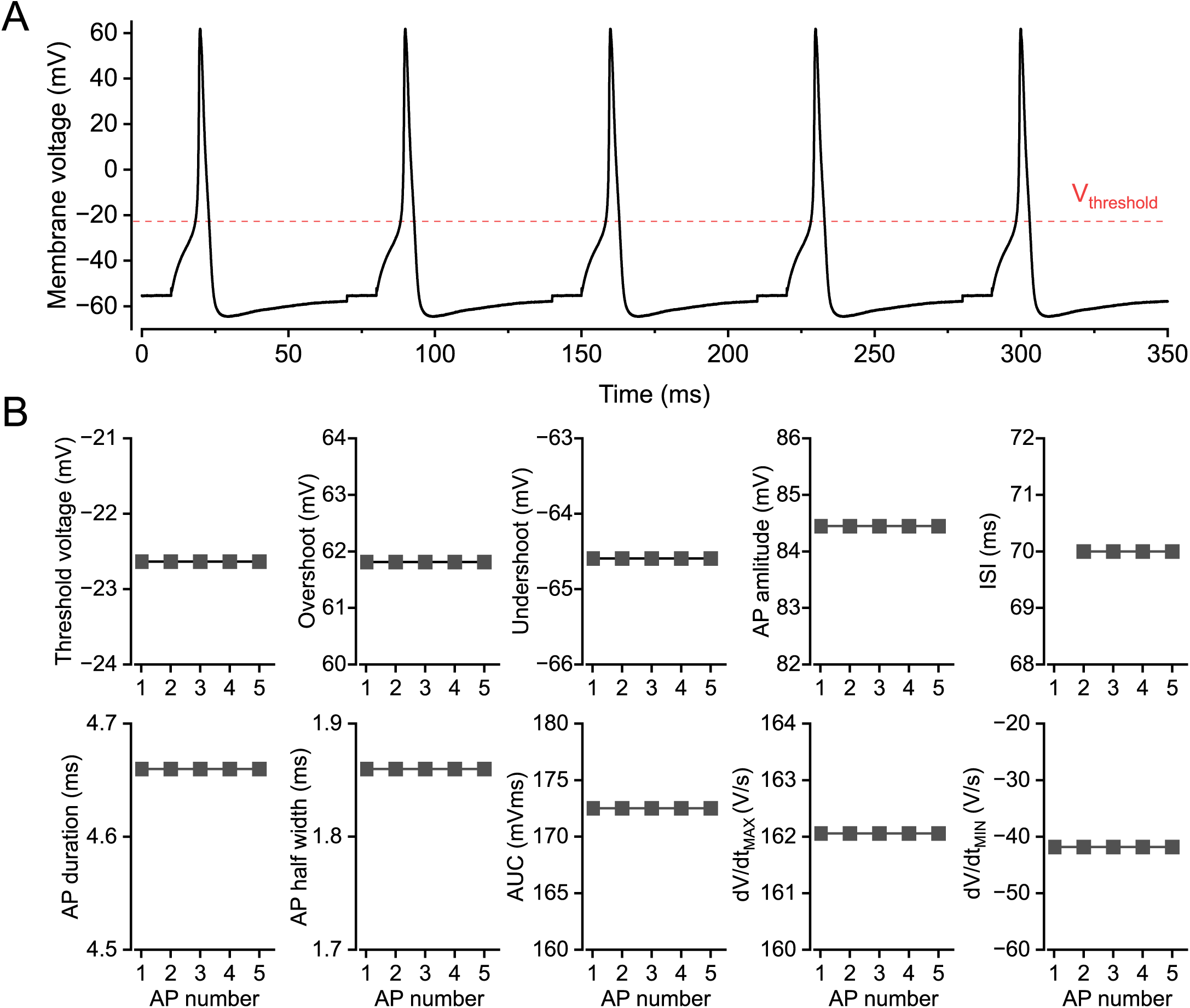
Stability of calculations performed on identical copied APs. Repeated analysis of identical copied AP waveforms (A) demonstrates stability of APpar parameter extraction (B). Threshold voltage, overshoot, undershoot, AP amplitude, ISI, AP duration, AP half-width, AP area above threshold, dV/dt_MAX_, and dV/dt_MIN_ remain stable across AP number when the input waveform is identical.

**Figure 6.**
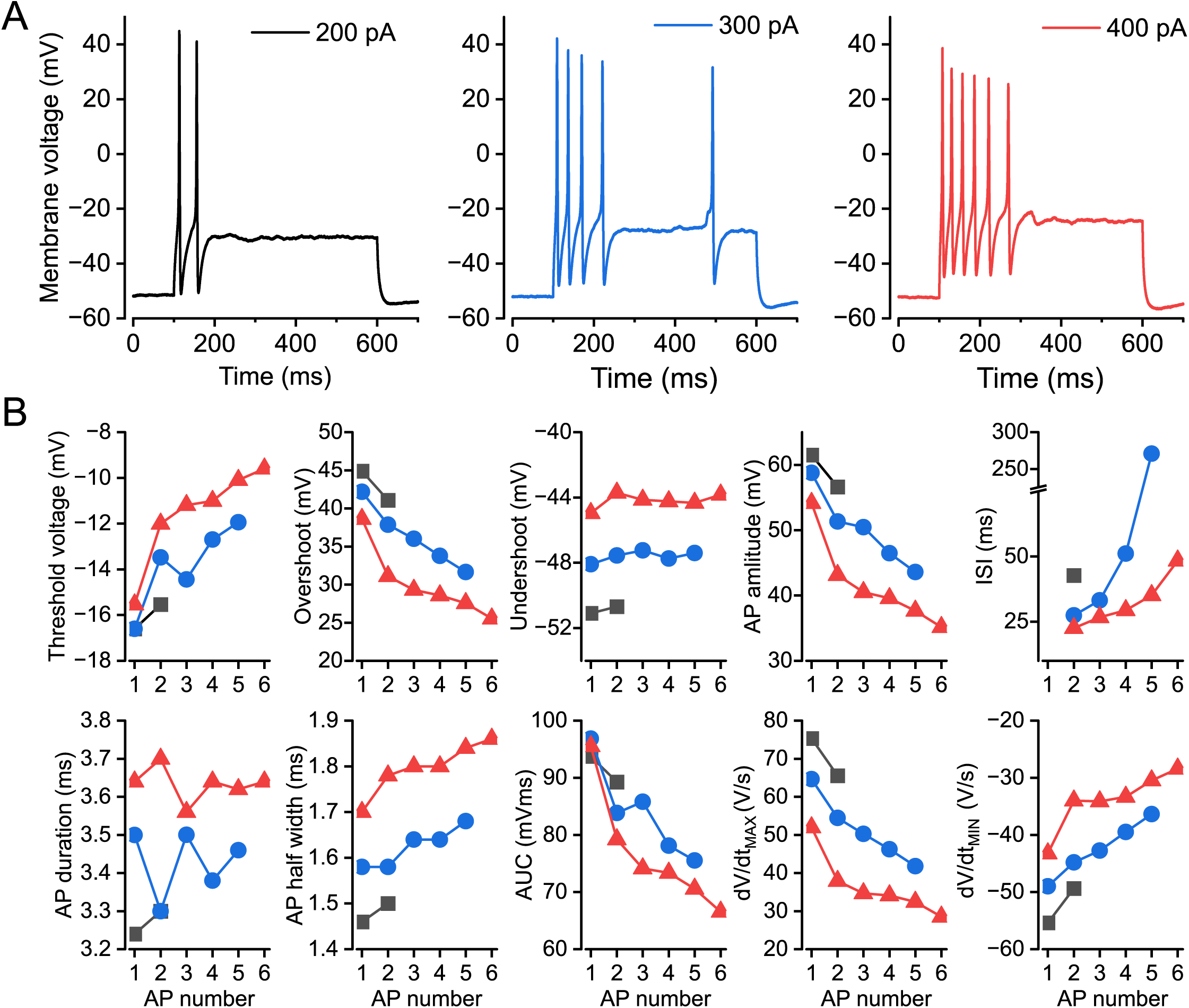
Representative recordings from a small DRG neuron. Representative current-evoked AP trains recorded from a small DRG neuron at different depolarizing current injections (A). APpar extracts AP-by-AP parameter trajectories including threshold voltage, overshoot, undershoot, AP amplitude, ISI, AP duration, AP half-width, AP area above threshold, dV/dt_MAX_, and dV/dt_MIN_(B).

**Figure 7.**
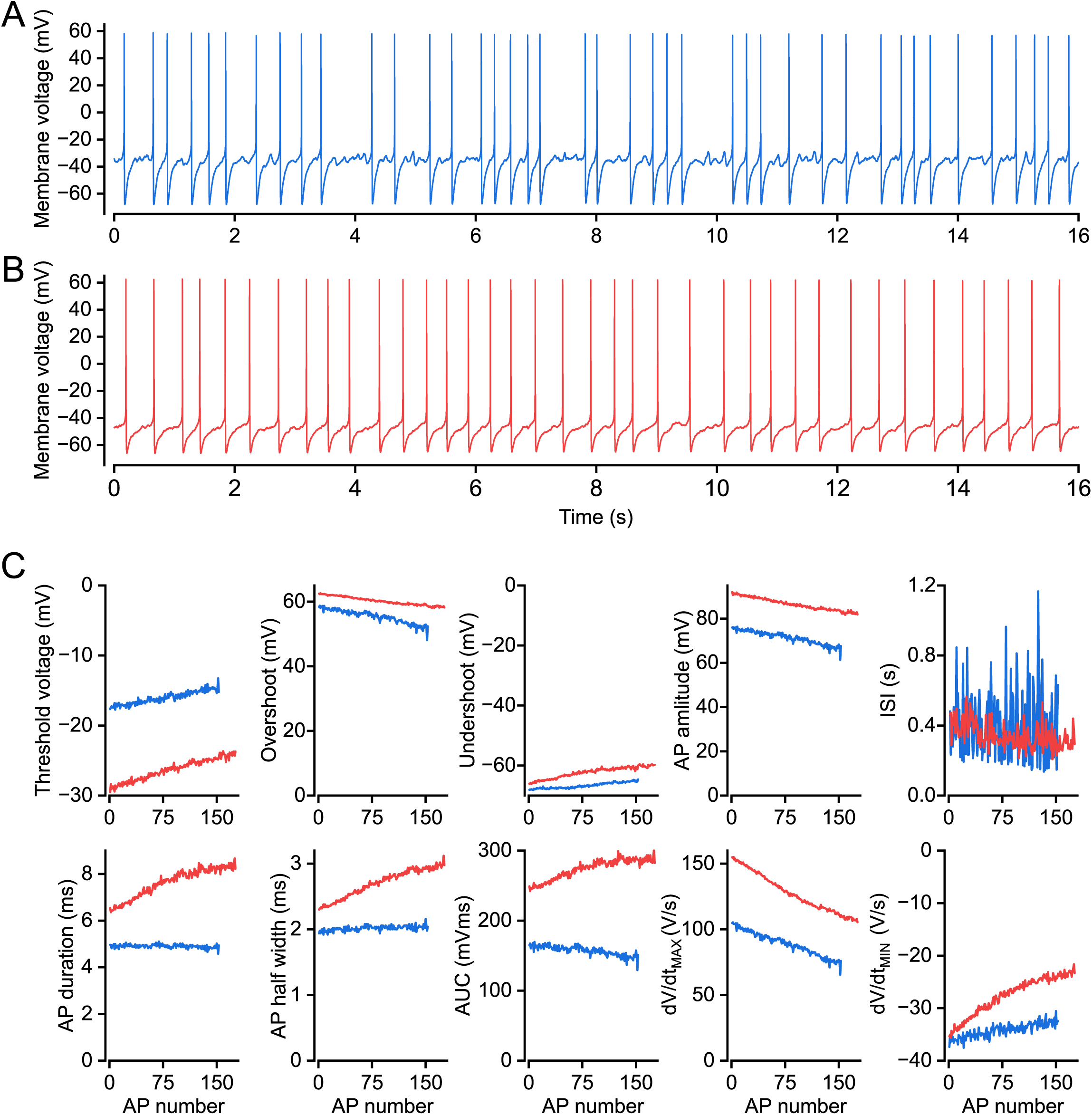
Representative recordings of spontaneous APs in two small DRG neurons. Long-duration spontaneous AP recordings from two small DRG neurons (A, B). The first 16 s are shown in the top panels (200 pA, 300 pA and 400 pA denote respective stimulus strength), while AP parameters were calculated from 60 s recordings. APpar quantifies dynamic changes in threshold voltage, overshoot, undershoot, AP amplitude, ISI, AP duration, half-width, AP area above threshold, dV/dt_MAX_, and dV/dt_MIN_ across spontaneous firing sequences (B). The color-code is the same across all corresponding panels and graphs.

### Software Availability

APpar software was developed and tested on Windows 10 and Windows 11 systems for use under OriginLab LabTalk and Origin-Python environments in Origin 2022b and OriginPro 2025. The source code and template Origin files will be provided upon paper acceptance as Extended Data in a single ZIP archive. All source code for APpar will be available upon paper acceptance on GitHub at http://github.com/dmytrovasylyev/APpar. APpar is released under the GNU General Public License (GPL) Version 3.

## Results

### APpar integrates AP analysis into an Origin/OriginPro electrophysiology workflow

APpar is designed to operate within the data-analysis environment already used by many electrophysiologists. Figure 1 shows the APpar workflow in Origin/OriginPro. The source worksheet contains time and voltage recordings. APpar assumes that input and output time values are in seconds and membrane voltage values are in Volts. A settings dialog defines trace number, RMP window, AP search window, derivative threshold, smoothing, overshoot and duration filters, minimal inter-AP interval, and minimum threshold-duration criterion. After execution, APpar writes derivative traces and parameter tables, allowing the user to inspect both numerical output and the derivative traces used for detection.

This workflow was designed to minimize manual steps while preserving user control over physiologically meaningful criteria. The dV/dt threshold remains user-defined because threshold criteria may need to differ across recording types, sampling frequencies, noise levels, recording temperature, and neuronal populations. By writing derivative traces into the source worksheet, APpar allows direct inspection of the relationship between voltage waveform, derivative trace, and detected threshold points.

### Algorithmic structure of APpar

Figure 2 summarizes the operational diagram and AP detection algorithm. APpar begins by reading the active (after activating by clicking on it) Origin worksheet, obtaining time from column A and voltage traces from subsequent columns. It then calculates raw dV/dt using a central-difference approximation and smooths the derivative using adjacent averaging. The software scans the user-defined AP search interval for positive dV/dt threshold crossings. Candidate APs are evaluated using inter-AP interval, overshoot, duration, and minimum derivative-duration criteria.

After the initial forward threshold crossing is found, APpar applies TRUE-threshold validation. This validation searches from AP overshoot backward to the closest preceding local dV/dt maximum and then further backward to the user-defined derivative threshold crossing.

The sample after this backward crossing is used as the validated threshold for all subsequent calculations. The output structure includes settings, AP-by-AP measurements, derivative values, and threshold-validation values.

### APpar formalizes definitions of AP waveform parameters and validates AP threshold determination

Figure 3 illustrates the core AP parameters calculated by the software. These include RMP, threshold voltage, overshoot, undershoot, AP amplitude, half amplitude, AP duration, half-width, dV/dt_MAX_, and dV/dt_MIN_. The central organizing feature is the validated threshold point. Because AP amplitude is calculated as overshoot minus threshold voltage, any change in threshold definition can change amplitude. Similarly, AP duration and AP area above threshold depend on the threshold reference voltage.

This is why APpar reports threshold-related information explicitly. The software does not simply output AP amplitude as a peak value or RMP-to-peak excursion; it defines amplitude relative to AP onset. This is consistent with Harty and Waxman (2007) (Patrick Harty and Waxman, 2007), who measured AP amplitude as the difference between peak and AP onset, with AP onset defined by the derivative criterion, and other studies (Prescott et al., 2008; Yi et al., 2015). This convention is particularly appropriate for DRG neuron studies because sodium channel availability and threshold mechanisms can shift independently of resting membrane potential.

The lower panel of Figure 3 illustrates AP threshold determination and TRUE-threshold validation. A fixed dV/dt criterion detects the first point at which the AP rising phase exceeds the selected derivative threshold. However, derivative traces can include small early peaks due to stimulation artifacts, noise, or pre-threshold fluctuations. APpar therefore uses the initial crossing as a candidate rather than a final endpoint.

The TRUE-threshold validation algorithm uses the structure of the AP rising phase. It first identifies AP overshoot, then searches backward to the nearest earlier local dV/dt maximum, which anchors the main AP upstroke. Blue arrows illustrate the evolution of the algorithmic search process. It then continues backward until the derivative falls below the user-defined criterion. The next point is treated as the TRUE threshold. This procedure retains the physiological interpretability of a derivative threshold while reducing the chance that an isolated earlier fluctuation defines the threshold.

The algorithm is transparent because both forward and TRUE thresholds are reported. Users can therefore evaluate whether threshold validation made substantial changes in a given dataset and can adjust smoothing or threshold criteria if the derivative trace is too noisy.

### Rheobase, peri-threshold region, and AP threshold voltage

Figure 4 illustrates rheobase determination and peri-threshold membrane behavior in a representative small DRG neuron. Current injection of 460 pA in this example cell did not evoke an action potential, whereas 470 pA evoked an AP, defining rheobase under these recording conditions. The lower panel highlights the peri-threshold region associated with AP initiation and threshold voltage determination.

These recordings demonstrate how relatively small changes in depolarizing current near rheobase can determine whether regenerative spike initiation occurs. The peri-threshold region is especially important for studies of excitability because it reflects the balance between inward and outward membrane currents immediately preceding AP initiation (Prescott et al., 2008; Yi et al., 2015). APpar enables extraction of threshold voltage and related waveform parameters from this transition region while preserving derivative-based threshold definitions.

### Stability of parameter extraction from identical copied APs

Figure 5 tests APpar stability using identical copied APs. The top trace shows repeated identical AP waveforms. The lower panels show extracted values for threshold voltage, overshoot, undershoot, AP amplitude, interspike interval (ISI, defined as the threshold-to-threshold timing between two adjacent APs and calculated post hoc by the user), AP duration, AP half-width, AP area above threshold, dV/dt_MAX_, and dV/dt_MIN_ across AP number.

The parameters remained stable across repeated copied APs, demonstrating that APpar produces reproducible measurements when the input waveform is identical. This validation is important because it separates analytical variability from biological variability. If repeated identical APs yielded drifting threshold, amplitude, or width values, the algorithm would not be suitable for detecting biological changes across repetitive firing. The observed stability supports the use of APpar for sequence analysis of AP trains.

### APpar quantifies parameter dynamics during current-evoked repetitive firing

Figure 6 shows representative current-evoked recordings from a small DRG neuron. APpar detected APs during responses to different current injection amplitudes and extracted multiple waveform parameters across AP number. The plots demonstrate that APpar can quantify not only spike count and ISI but also progressive changes in threshold voltage, overshoot, undershoot, AP amplitude, AP duration, half-width, AP area, dV/dt_MAX_, and dV/dt_MIN_.

These measurements are relevant because repetitive firing is shaped by cumulative inactivation and recovery of sodium channels, activation of potassium currents, changes in membrane resistance, and activity-dependent adaptation. A single parameter such as AP number is therefore insufficient to describe the full excitability phenotype. APpar provides a richer output that allows the user to determine whether firing changes are associated with threshold shifts, spike broadening, reduced upstroke speed, altered repolarization, or changes in afterhyperpolarization.

### APpar analyzes long-duration spontaneous AP recordings

Figure 7 shows spontaneous AP recordings from two small DRG neurons. The top panels display the first 16 s of recordings, while AP parameters were calculated from 60 s recordings. APpar extracted parameter trajectories across more than 150 APs. The output shows dynamic changes in threshold voltage, overshoot, undershoot, AP amplitude, ISI, AP duration, half-width, AP area, dV/dt_MAX_, and dV/dt_MIN_.

Spontaneous AP recordings are challenging because AP intervals are not imposed by a stimulus protocol, baseline can drift, and waveform properties can evolve over time. APpar handles these recordings by applying derivative-based detection and AP-specific parameter extraction to each detected event. The resulting parameter trajectories reveal patterns that would be difficult and time-consuming to obtain manually.

## Discussion

APpar provides an automated, transparent, and Origin-compatible software workflow for AP analysis in neuronal electrophysiology. The software extracts a broad set of waveform parameters and explicitly defines the threshold-dependent calculations that often vary across laboratories. Its central algorithmic contribution is TRUE-threshold validation, which preserves the practical value of derivative-based threshold detection while reducing susceptibility to early derivative fluctuations.

The decision to define AP amplitude as overshoot minus threshold voltage is important. Electrophysiologists refer to AP amplitude in different ways, including RMP-to-peak amplitude, threshold-to-peak amplitude, or peak voltage alone. These are not interchangeable. RMP-to-peak amplitude includes baseline membrane polarization, whereas threshold-to-peak amplitude isolates the voltage excursion after AP initiation. Harty and Waxman (2007) (Patrick Harty and Waxman, 2007) and others (Prescott et al., 2008; Yi et al., 2015) used the latter logic by defining AP amplitude as the difference between peak and AP onset, with AP onset defined by a dV/dt criterion. APpar follows this operational logic and makes the definition explicit. Because alternative definitions are also commonly used in the literature, APpar reports all underlying voltage parameters for each analyzed AP to avoid bias toward any single convention for AP amplitude determination, thereby enabling users to apply their preferred definition during downstream analysis.

This matters for neurons such as DRG neurons because sodium channel availability, threshold voltage, and AP peak can shift independently. For example, (Patrick Harty and Waxman, 2007) showed that AP amplitude in small DRG neurons decreases as membrane potential is depolarized and that this relationship reflects contributions of TTX-sensitive sodium channels and Nav1.8. More recent dynamic clamp studies further emphasize the importance of channel interactions near threshold. It was demonstrated that Nav1.8 and Nav1.7 interact strongly close to AP threshold and that partial Nav1.8 subtraction can reverse hyperexcitability in a dynamic clamp model of nociceptor hyperexcitability (Vasylyev et al., 2024). However, this effect is context-dependent and, in a subpopulation of nociceptors, is modest due to intrinsic limits on the functional effect of Nav1.8 abrogation. These and other studies (Xie et al., 2022; Xie et al., 2024; Stewart et al., 2025; Fujita et al., 2026) underscore the need for analysis tools that treat threshold not as a casual cursor placement but as a defined measurement tied to ion channel function.

APpar also formalizes parameters beyond threshold and amplitude. AP duration, AP half-width, AP width at 0 mV, and AP area above threshold provide complementary measures of spike waveform. dV/dt_MAX_ and dV/dt_MIN_ provide derivative-based readouts of the fastest depolarizing and repolarizing phases. Undershoot and undershoot latency capture the afterhyperpolarizing phase. ISI and AP number capture firing organization. Together, these parameters support multidimensional analysis of excitability rather than reliance on a single summary variable.

The validation figures demonstrate several practical strengths. Identical copied APs produced stable parameter values, indicating that the analysis does not introduce artificial drift. Analysis of AP firing in DRG neuron in response to long-duration current pulses showed that APpar can analyze repetitive firing and extract AP-by-AP dynamics. Recordings of spontaneous AP firing showed that APpar can analyze long-duration datasets of irregular AP firing. The software therefore supports both stimulus-evoked and spontaneous activity workflows. The default settings were optimized for APs recorded from small DRG neurons at room temperature (23 °C); however, the user-friendly parameter settings interface implies broad utility of the program across a wide range of neuronal types and, perhaps, non-neuronal cells.

The Origin-Python implementation is also useful from a practical laboratory perspective. Many electrophysiologists use Origin or OriginPro for visualization, fitting, and figure preparation. By embedding Python analysis inside an Origin workflow, APpar allows users to retain familiar data handling while gaining speed and more flexible code structure. This is especially useful for laboratories that need analysis transparency but do not want to abandon existing Origin-based workflows.

Several limitations remain. First, threshold is still criterion-dependent. APpar improves consistency by making the criterion explicit and by validating threshold placement, but the selected dV/dt criterion and smoothing window remain user decisions. Second, derivative-based methods are sensitive to sampling rate and noise. Users should inspect derivative traces and adjust smoothing appropriately, particularly to avoid altering derivative kinetics or artificially reducing dV/dt_MAX_ or dV/dt_MIN_. Third, the present validation is based on representative recordings from small DRG neurons; broader validation across neuronal classes, sampling frequencies, and recording conditions would further establish generalizability. Fourth, APpar currently focuses on AP waveform extraction rather than automated classification of firing phenotypes or burst patterns.

Future development may include additional threshold modes, such as second derivative or hybrid derivative/curvature methods, automated baseline drift correction, burst analysis, refractory-cycle analysis, and direct import from additional electrophysiology file formats. A graphical threshold-quality report could also help users identify recordings where derivative noise requires manual review.

In summary, APpar provides a practical and reproducible tool for AP waveform analysis in neuronal electrophysiology. By combining automated parameter extraction, explicit threshold definitions, TRUE-threshold validation, and Origin-compatible output, APpar can reduce manual analysis burden and improve reproducibility in studies of neuron excitability and ion channel function.

## Author Contributions

D.V.V. conceived the software, designed and developed it, performed the research, analyzed and interpreted the data, and wrote the manuscript. S.G.W. interpreted the data, acquired funding, reviewed and edited the manuscript.

## Conflict of Interest

The author declares no competing financial interests.

## Acknowledgements

This work was supported in part by Grant RX002999-01, Department of Veterans Affairs and through funding from the Paralyzed Veterans of America, Erythromelalgia Association, Crenshaw Fund, and the Bridget Flaherty Endowment.

## Notes

http://github.com/dmytrovasylyev/APpar

### Competing Interest Statement

The authors have declared no competing interest.

