## Supplementary material for "APpar: automated action potential parameter analysis software for reproducible electrophysiological measurements in neurons": 0_ReadMe Installation APpar on LabTalk-Python.txt

APpar.ogs
APpar_2026-05-11_v1.0_python is based on APpar_2026-05-11_v1.0.ogs
APpar_2026-05-11_v1.0_python fixed_for_Origin2022b

ATTN: all input values of APs (time and voltage) must be in seconds and Volts

==================================================================================================This part needs to be done only once per PC life
// OriginPro2025 or Origin2022b or later:
// Open from Origin menu: Connectivity - Python Packages
// Install the following packages:
numpy
scipy
pandas

// Open from Origin menu: Connectivity - Python Console
import sys
//ENTER
print(sys.executable)
//ENTER
import subprocess
//ENTER
subprocess.check_call([sys.executable, "-m", "pip", "install", "numpy", "scipy", "pandas"])
//ENTER

//re-start Origin
==================================================================================================This part needs to be done only once per PC life
==================================================================================================


=====================================================================================================This part needs to be done for every APpar or APsim program (APsim17, APsim18, etc)
//APpar.ogs
// Wrapper to run APpar.py from Origin with the same settings dialog style.
// Attn: in the file APpar.ogs replace dv93 with your username to update Python script location on your PC:
string pyFile$ = "C:\\Users\\dv93\\Documents\\OriginLab\\User Files\\APpar.py";

//APpar.ogs and APpar.py
// Attn: Place APpar.ogs and APpar.py files into the following directory in your PC (replace dv93 with your username):
C:\Users\dv93\Documents\OriginLab\User Files

//re-start Origin

=====================================================================================================This part needs to be done for every APpar or APsim program (APsim17, APsim18, etc)

=====================================================================================================Run the program
// Make sure Book/spreadsheet with the AP traces to be calculated is active (click on it)
// Open LabTalk Command Window from Origin menu: Window - Command Window or (Alt+3)
//Type APpar in the LabTalk Command Window and hit ENTER
=====================================================================================================Run the program
