## Supplementary material for "APpar: automated action potential parameter analysis software for reproducible electrophysiological measurements in neurons": APpar.py

```
# ============================================================================
# APpar - Action Potential Parameters Analyzer
##
# Developed in 2026 by Dmytro Vasylyev
# Yale University
#
# Copyright (c) 2026 Dmytro Vasylyev.
# All rights reserved.
#
# Distribution, licensing, and commercialization rights may be subject to
# institutional review and approval requirements.
#
# Contact:
#  
#
# APpar.py
# Python conversion of APparLT.ogs / LabTalk

from __future__ import annotations

from dataclasses import dataclass
from typing import Any, List, Optional, Tuple

import numpy as np
import pandas as pd

try:
    import originpro as op
except Exception as exc:  # pragma: no cover - meant for Origin
    op = None
    _IMPORT_ERROR = exc
else:
    _IMPORT_ERROR = None


PARAMETER_NAMES = [
    "Trace number",
    "Source voltage column",
    "RMP start time (s)",
    "RMP end time (s)",
    "AP search start time (s)",
    "AP search end time (s)",
    "dV/dt threshold (V/s)",
    "dV/dt filter number of points",
    "Minimal AP overshoot (V)",
    "Minimal AP duration (s)",
    "Minimal inter-AP interval (s)",
    "Minimum dV/dt threshold duration (s)",
    "RMP (V)",
    "AP number",
    "Threshold row",
    "Threshold time (s)",
    "Threshold voltage (V)",
    "dV/dt at threshold (V/s)",
    "Overshoot (V)",
    "Overshoot time (s)",
    "Undershoot (V)",
    "Undershoot time (s)",
    "Undershoot latency from threshold (s)",
    "AP amplitude (V)",
    "AP half amplitude (V)",
    "AP half amplitude time (s)",
    "AP rise time (s)",
    "AP rise start time (s)",
    "AP rise end time (s)",
    "AP decay time (s)",
    "AP duration (s)",
    "AP duration start (s)",
    "AP duration end (s)",
    "AP half-width (s)",
    "AP half-width start (s)",
    "AP half-width end (s)",
    "AP width at 0 mV (s)",
    "AP width at 0 mV start (s)",
    "AP width at 0 mV end (s)",
    "AP area above threshold ((V-Vthr)*s)",
    "dV/dt max (V/s)",
    "dV/dt max time (s)",
    "dV/dt min (V/s)",
    "dV/dt min time (s)",
    "Interspike interval threshold-to-threshold (s)",
    "Number of APs detected",
    "Forward threshold row",
    "Forward threshold time (s)",
    "TRUE threshold row",
    "TRUE threshold time (s)",
    "Threshold source",
    "Forward vs TRUE row difference",
]


@dataclass
class Settings:
    n_traces: int = 1
    rmp_start_time: float = 0.004
    rmp_end_time: float = 0.009
    ap_start_time: float = 0.01
    ap_end_time: Optional[float] = None
    dvdt_crit: float = 8.0
    dvdt_smooth_points: int = 6
    min_overshoot: float = 0.0
    min_ap_duration: float = 0.001
    min_isi: float = 0.01
    min_dvdt_threshold_duration: float = 0.0002
    write_pause_sec: float = 0.025
    # Fixed voltage margin used for AP validation, in volts.
    # The AP overshoot must exceed the validated/rising threshold Vm by this
    # amount, and the peak must then be confirmed by a post-peak Vm fall of at
    # least this amount. This additional filter rejects noisy dV/dt threshold
    # crossings/stimulus artifacts and prevents equal/near-equal Vm samples near
    # threshold from being accepted as APs, especially if the user sets minimal
    # AP duration to 0. The falling dV/dt crossing Vm is NOT used as a required
    # voltage reference because it can be very close to the AP peak.
    overshoot_crossing_margin: float = 0.003


def _origin_type(msg: str) -> None:
    if op is not None:
        try:
            op.lt_exec(f'type "{msg}";')
            return
        except Exception:
            pass
    print(msg)


def _to_numeric_array(values: Any) -> np.ndarray:
    return pd.to_numeric(pd.Series(values), errors="coerce").to_numpy(dtype=float)


def _moving_average_same(y: np.ndarray, npts: int) -> np.ndarray:
    npts = int(max(1, npts))
    if npts <= 1 or len(y) == 0:
        return y.copy()
    out = np.empty_like(y, dtype=float)
    half_left = (npts - 1) // 2
    half_right = npts // 2
    for i in range(len(y)):
        a = max(0, i - half_left)
        b = min(len(y), i + half_right + 1)
        out[i] = np.nanmean(y[a:b])
    return out


def _derivative_dvdt(time: np.ndarray, voltage: np.ndarray) -> np.ndarray:
    if len(time) < 2:
        return np.zeros_like(voltage, dtype=float)
    dvdt = np.empty_like(voltage, dtype=float)
    dvdt[0] = (voltage[1] - voltage[0]) / (time[1] - time[0])
    dvdt[-1] = (voltage[-1] - voltage[-2]) / (time[-1] - time[-2])
    dt = time[2:] - time[:-2]
    dv = voltage[2:] - voltage[:-2]
    dvdt[1:-1] = dv / dt
    return dvdt


def _first_index_ge(x: np.ndarray, value: float) -> int:
    matches = np.where(x >= value)[0]
    return int(matches[0]) if len(matches) else 0


def _last_index_le(x: np.ndarray, value: float) -> int:
    matches = np.where(x <= value)[0]
    return int(matches[-1]) if len(matches) else len(x) - 1


def analyze_trace(time: np.ndarray, voltage: np.ndarray, dvdt_sm: np.ndarray, trace_num: int,
                  trace_col_1based: int, settings: Settings) -> Tuple[pd.DataFrame, int]:
    t_end = settings.ap_end_time if settings.ap_end_time is not None else float(time[-1])
    i_start = _first_index_ge(time, settings.ap_start_time)
    i_end = _last_index_le(time, t_end)

    rmp_mask = (time >= settings.rmp_start_time) & (time <= settings.rmp_end_time)
    rmp = float(np.nanmean(voltage[rmp_mask])) if np.any(rmp_mask) else 0.0

    rows = {name: [np.nan] for name in PARAMETER_NAMES}
    settings_values = [
        trace_num, trace_col_1based, settings.rmp_start_time, settings.rmp_end_time,
        settings.ap_start_time, t_end, settings.dvdt_crit, settings.dvdt_smooth_points,
        settings.min_overshoot, settings.min_ap_duration, settings.min_isi,
        settings.min_dvdt_threshold_duration, rmp,
    ]
    for idx, value in enumerate(settings_values):
        rows[PARAMETER_NAMES[idx]][0] = value

    ap_records: List[dict] = []
    ap_count = 0
    last_ap_time = -999999.0
    i = i_start + 1

    while i <= i_end:
        if dvdt_sm[i] >= settings.dvdt_crit and dvdt_sm[i - 1] < settings.dvdt_crit:
            i_forward_thr = i
            t_forward_thr = float(time[i_forward_thr])
            v_forward_thr = float(voltage[i_forward_thr])

            if (t_forward_thr - last_ap_time) >= settings.min_isi:
                # Preliminary overshoot search for candidate validation.
                # The search is not allowed to terminate until BOTH conditions
                # are satisfied: (1) the user-defined minimal AP-duration window
                # has elapsed, and (2) a true local maximum has been confirmed by
                # a post-peak Vm fall of at least 0.003 V. This rejects false
                # overshoots caused by small Vm noise or flat/equal adjacent Vm
                # samples, including when min_ap_duration is set to 0.
                i_original_peak = i_forward_thr
                over_val = float(voltage[i_forward_thr])
                over_time = float(time[i_forward_thr])
                t_prelim_min_end = t_forward_thr + settings.min_ap_duration
                prelim_peak_confirmed = False
                for j in range(i_forward_thr, i_end + 1):
                    if voltage[j] > over_val:
                        over_val = float(voltage[j])
                        i_original_peak = j
                        over_time = float(time[j])
                        prelim_peak_confirmed = False
                    if j > i_original_peak and voltage[j] <= over_val - settings.overshoot_crossing_margin:
                        prelim_peak_confirmed = True
                    if time[j] >= t_prelim_min_end and prelim_peak_confirmed:
                        break

                if not prelim_peak_confirmed:
                    i += 1
                    continue

                # TRUE threshold: start from dV/dtMAX during the AP upstroke,
                # then continue backward until dV/dt drops below user criterion.
                # This avoids anchoring the search to AP overshoot voltage.
                i_dvdt_max_anchor = i_forward_thr
                max_dvdt_val = float(dvdt_sm[i_forward_thr])
                for j in range(i_forward_thr, i_original_peak + 1):
                    if dvdt_sm[j] > max_dvdt_val:
                        max_dvdt_val = float(dvdt_sm[j])
                        i_dvdt_max_anchor = j

                i_true_thr = i_forward_thr
                for j in range(i_dvdt_max_anchor, i_start - 1, -1):
                    if dvdt_sm[j] < settings.dvdt_crit:
                        i_true_thr = j + 1
                        break

                i_thr = i_true_thr
                t_thr = float(time[i_thr])
                v_thr = float(voltage[i_thr])
                dvdt_thr = float(dvdt_sm[i_thr])
                t_true_thr = float(time[i_true_thr])
                threshold_row_difference = abs((i_true_thr + 1) - (i_forward_thr + 1))

                # Recalculate AP peak from TRUE threshold. The overshoot-search
                # cycle is not allowed to terminate until BOTH conditions are
                # satisfied: (1) the user-defined minimal AP-duration window has
                # elapsed, and (2) the candidate local maximum is confirmed by a
                # post-peak Vm fall of at least 0.003 V. This prevents false
                # overshoots caused by Vm noise or flat/equal adjacent samples.
                i_peak = i_thr
                over_val = float(voltage[i_thr])
                over_time = float(time[i_thr])
                t_min_ap_end = t_thr + settings.min_ap_duration
                peak_confirmed = False
                for j in range(i_thr, i_end + 1):
                    if voltage[j] > over_val:
                        over_val = float(voltage[j])
                        i_peak = j
                        over_time = float(time[j])
                        peak_confirmed = False
                    if j > i_peak and voltage[j] <= over_val - settings.overshoot_crossing_margin:
                        peak_confirmed = True
                    if time[j] >= t_min_ap_end and peak_confirmed:
                        break

                # dV/dt must return below criterion after the AP overshoot before
                # the voltage-return AP-duration endpoint is accepted.
                i_dvdt_thresh_end = i_peak
                for j in range(i_peak, i_end + 1):
                    if dvdt_sm[j] < settings.dvdt_crit:
                        i_dvdt_thresh_end = j
                        break
                    i_dvdt_thresh_end = j
                dvdt_duration = float(time[i_dvdt_thresh_end] - t_thr)
                v_dvdt_fall_cross = float(voltage[i_dvdt_thresh_end])

                i_dur_end = -1
                for j in range(max(i_peak + 1, i_dvdt_thresh_end), i_end + 1):
                    if voltage[j] <= v_thr:
                        i_dur_end = j
                        break

                if i_dur_end > 0:
                    ap_duration_start = t_thr
                    ap_duration_end = float(time[i_dur_end])
                    ap_duration = ap_duration_end - ap_duration_start

                    pass_ap = True
                    if over_val < settings.min_overshoot:
                        pass_ap = False

                    # Additional noise/artifact filter: overshoot must be at least
                    # 0.003 V above the validated/rising threshold Vm. The falling
                    # dV/dt crossing Vm is NOT used here because it can be very close
                    # to the AP peak and would falsely reject real narrow APs. A true
                    # local maximum is instead confirmed by the post-peak Vm fall of
                    # at least 0.003 V, represented by peak_confirmed.
                    if over_val < v_thr + settings.overshoot_crossing_margin:
                        pass_ap = False
                    if not peak_confirmed:
                        pass_ap = False

                    if ap_duration < settings.min_ap_duration:
                        pass_ap = False
                    if dvdt_duration < settings.min_dvdt_threshold_duration:
                        pass_ap = False

                    if pass_ap:
                        interspike_interval = np.nan if ap_count == 0 else t_thr - last_ap_time
                        ap_count += 1
                        last_ap_time = t_thr

                        i_under_end = i_end
                        for j in range(i_peak + 1, i_end + 1):
                            if dvdt_sm[j] >= settings.dvdt_crit and dvdt_sm[j - 1] < settings.dvdt_crit:
                                i_under_end = j - 1
                                break

                        i_under = i_peak
                        under_val = float(voltage[i_peak])
                        for j in range(i_peak, i_under_end + 1):
                            if voltage[j] < under_val:
                                under_val = float(voltage[j])
                                i_under = j

                        under_time = float(time[i_under])
                        under_latency = under_time - t_thr
                        ap_rise_start = t_thr
                        ap_rise_end = over_time
                        ap_rise_time = ap_rise_end - ap_rise_start
                        ap_decay_time = ap_duration_end - over_time

                        ap_area = 0.0
                        for j in range(i_thr, i_dur_end):
                            dt = float(time[j + 1] - time[j])
                            v1 = float(voltage[j] - v_thr)
                            v2 = float(voltage[j + 1] - v_thr)
                            ap_area += 0.5 * (v1 + v2) * dt

                        ap_amplitude = over_val - v_thr
                        ap_half_amplitude = v_thr + 0.5 * (over_val - v_thr)

                        i_half_start = next((j for j in range(i_thr, i_peak + 1)
                                             if voltage[j] >= ap_half_amplitude), None)
                        ap_half_amplitude_time = float(time[i_half_start]) if i_half_start is not None else 0.0

                        i_half_end = next((j for j in range(i_peak, i_dur_end + 1)
                                           if voltage[j] <= ap_half_amplitude), None)
                        if i_half_start is not None and i_half_end is not None:
                            ap_half_width_start = float(time[i_half_start])
                            ap_half_width_end = float(time[i_half_end])
                            ap_half_width = ap_half_width_end - ap_half_width_start
                        else:
                            ap_half_width = 0.0
                            ap_half_width_start = 0.0
                            ap_half_width_end = 0.0

                        i_zero_up = next((j for j in range(i_thr, i_peak + 1) if voltage[j] >= 0), None)
                        i_zero_down = next((j for j in range(i_peak, i_dur_end + 1) if voltage[j] <= 0), None)
                        if i_zero_up is not None and i_zero_down is not None:
                            ap_width0_start = float(time[i_zero_up])
                            ap_width0_end = float(time[i_zero_down])
                            ap_width0 = ap_width0_end - ap_width0_start
                        else:
                            ap_width0 = 0.0
                            ap_width0_start = 0.0
                            ap_width0_end = 0.0

                        seg = dvdt_sm[i_thr:i_dur_end + 1]
                        rel_max = int(np.nanargmax(seg))
                        rel_min = int(np.nanargmin(seg))
                        i_dvdt_max = i_thr + rel_max
                        i_dvdt_min = i_thr + rel_min

                        ap_records.append({
                            "AP number": ap_count,
                            "Threshold row": i_thr + 1,
                            "Threshold time (s)": t_thr,
                            "Threshold voltage (V)": v_thr,
                            "dV/dt at threshold (V/s)": dvdt_thr,
                            "Overshoot (V)": over_val,
                            "Overshoot time (s)": over_time,
                            "Undershoot (V)": under_val,
                            "Undershoot time (s)": under_time,
                            "Undershoot latency from threshold (s)": under_latency,
                            "AP amplitude (V)": ap_amplitude,
                            "AP half amplitude (V)": ap_half_amplitude,
                            "AP half amplitude time (s)": ap_half_amplitude_time,
                            "AP rise time (s)": ap_rise_time,
                            "AP rise start time (s)": ap_rise_start,
                            "AP rise end time (s)": ap_rise_end,
                            "AP decay time (s)": ap_decay_time,
                            "AP duration (s)": ap_duration,
                            "AP duration start (s)": ap_duration_start,
                            "AP duration end (s)": ap_duration_end,
                            "AP half-width (s)": ap_half_width,
                            "AP half-width start (s)": ap_half_width_start,
                            "AP half-width end (s)": ap_half_width_end,
                            "AP width at 0 mV (s)": ap_width0,
                            "AP width at 0 mV start (s)": ap_width0_start,
                            "AP width at 0 mV end (s)": ap_width0_end,
                            "AP area above threshold ((V-Vthr)*s)": ap_area,
                            "dV/dt max (V/s)": float(dvdt_sm[i_dvdt_max]),
                            "dV/dt max time (s)": float(time[i_dvdt_max]),
                            "dV/dt min (V/s)": float(dvdt_sm[i_dvdt_min]),
                            "dV/dt min time (s)": float(time[i_dvdt_min]),
                            "Interspike interval threshold-to-threshold (s)": interspike_interval,
                            "Forward threshold row": i_forward_thr + 1,
                            "Forward threshold time (s)": t_forward_thr,
                            "TRUE threshold row": i_true_thr + 1,
                            "TRUE threshold time (s)": t_true_thr,
                            "Threshold source": "TRUE threshold used",
                            "Forward vs TRUE row difference": threshold_row_difference,
                        })
                        i = i_dur_end
        i += 1

    rows["Number of APs detected"][0] = ap_count
    result = pd.DataFrame({"Parameter": PARAMETER_NAMES, "Settings": [rows[name][0] for name in PARAMETER_NAMES]})
    for k, record in enumerate(ap_records, start=1):
        result[f"AP {k}"] = [record.get(name, np.nan) for name in PARAMETER_NAMES]
    return result, ap_count


def _read_active_worksheet() -> Tuple[Any, pd.DataFrame]:
    if op is None:
        raise RuntimeError(f"originpro could not be imported: {_IMPORT_ERROR}")

    wks = op.find_sheet()
    if wks is None:
        raise RuntimeError("No active worksheet found. Activate the source worksheet and run again.")

    # Origin 2022b can fail with wks.to_df() if several columns share the same
    # long name, for example many voltage columns named Vm. Read by position.
    ncols = int(wks.cols)
    data = {}
    max_len = 0
    for c in range(ncols):
        vals = wks.to_list(c)
        max_len = max(max_len, len(vals))
        data[f"Col{c + 1}"] = vals
    for key in data:
        if len(data[key]) < max_len:
            data[key] = data[key] + [np.nan] * (max_len - len(data[key]))
    df = pd.DataFrame(data)
    return wks, df


def _lt_value(name: str, default: Any, cast: Any) -> Any:
    if op is None:
        return default
    for func_name in ("lt_float", "lt_int", "get_lt_var"):
        func = getattr(op, func_name, None)
        if func is None:
            continue
        try:
            val = func(name)
            if val is not None:
                return cast(val)
        except Exception:
            pass
    return default


def settings_from_labtalk_or_defaults() -> Settings:
    s = Settings()
    s.n_traces = _lt_value("AP_NTRACES", s.n_traces, int)
    s.rmp_start_time = _lt_value("AP_RMP_START", s.rmp_start_time, float)
    s.rmp_end_time = _lt_value("AP_RMP_END", s.rmp_end_time, float)
    s.ap_start_time = _lt_value("AP_T_START", s.ap_start_time, float)
    ap_end = _lt_value("AP_T_END", np.nan, float)
    s.ap_end_time = None if np.isnan(ap_end) else ap_end
    s.dvdt_crit = _lt_value("AP_DVDT_CRIT", s.dvdt_crit, float)
    s.dvdt_smooth_points = _lt_value("AP_DVDT_SMOOTH_POINTS", s.dvdt_smooth_points, int)
    s.min_overshoot = _lt_value("AP_MIN_OVERSHOOT", s.min_overshoot, float)
    s.min_ap_duration = _lt_value("AP_MIN_AP_DURATION", s.min_ap_duration, float)
    s.min_isi = _lt_value("AP_MIN_ISI", s.min_isi, float)
    s.min_dvdt_threshold_duration = _lt_value("AP_MIN_DVDT_THRESHOLD_DURATION", s.min_dvdt_threshold_duration, float)
    s.write_pause_sec = _lt_value("AP_WRITE_PAUSE_SEC", s.write_pause_sec, float)
    return s


def _add_source_column(wks: Any, values: np.ndarray, lname: str) -> None:
    try:
        col_index = wks.cols
        wks.from_list(col_index, values.tolist(), lname=lname)
        return
    except Exception:
        pass
    try:
        wks.from_list(len(wks.to_df().columns), values.tolist(), lname)
        return
    except Exception:
        pass
    raise RuntimeError("Could not add/write a temporary derivative column to the source worksheet.")


def _new_result_book() -> Any:
    try:
        return op.new_book("w", lname="Book2")
    except Exception:
        try:
            return op.new_book(lname="Book2")
        except Exception:
            return op.new_book()


def _origin_safe_df(df: pd.DataFrame) -> pd.DataFrame:
    safe = df.copy()
    for col in safe.columns:
        dtype_name = str(safe[col].dtype).lower()
        if dtype_name in ("string", "stringdtype") or dtype_name.startswith("string[") or safe[col].dtype == object:
            vals = []
            for v in safe[col].tolist():
                if pd.isna(v):
                    vals.append("")
                else:
                    vals.append(str(v) if isinstance(v, str) else v)
            safe[col] = pd.Series(vals, dtype=object)
    return safe


def _write_df_to_wks(wks: Any, df: pd.DataFrame) -> None:
    safe = _origin_safe_df(df)
    try:
        wks.from_df(safe)
        return
    except Exception:
        for c, name in enumerate(safe.columns):
            wks.from_list(c, safe.iloc[:, c].tolist(), lname=str(name))
        return


def _write_result_sheet(book: Any, trace_num: int, df: pd.DataFrame) -> None:
    sheet_name = f"trace{trace_num}"
    try:
        if trace_num == 1:
            wks = book[0]
            try:
                wks.name = sheet_name
            except Exception:
                pass
        else:
            wks = book.add_sheet(sheet_name)
        _write_df_to_wks(wks, df)
        return
    except Exception:
        wks = op.new_sheet("w", lname=sheet_name)
        _write_df_to_wks(wks, df)


def run_ap_analysis(settings: Optional[Settings] = None) -> None:
    if settings is None:
        settings = Settings()

    wks, df = _read_active_worksheet()
    if df.shape[1] < 2:
        raise RuntimeError("Source worksheet must contain Col(A) time and at least Col(B) voltage.")

    time_all = _to_numeric_array(df.iloc[:, 0])
    valid_time = np.where(~np.isnan(time_all))[0]
    if len(valid_time) == 0:
        _origin_type("Error: no populated time values found in Col(A). Check that Col(A) contains the time data. Analysis stopped.")
        return

    n_data = int(valid_time[-1]) + 1
    time = time_all[:n_data]
    if settings.ap_end_time is None:
        settings.ap_end_time = float(time[-1])

    input_ncols = df.shape[1]
    result_book = _new_result_book()

    for trace_num in range(1, settings.n_traces + 1):
        trace_col_zero = trace_num  # trace1 is Col(B)
        trace_col_1based = trace_col_zero + 1
        if trace_col_zero >= input_ncols:
            _origin_type(f"Note: Trace {trace_num} was requested, but source column {trace_col_1based} does not exist. Analysis stopped before this trace.")
            break

        voltage = _to_numeric_array(df.iloc[:n_data, trace_col_zero])
        dvdt_raw = _derivative_dvdt(time, voltage)
        dvdt_sm = _moving_average_same(dvdt_raw, settings.dvdt_smooth_points)

        _add_source_column(wks, dvdt_raw, f"dV/dt_raw_trace{trace_num}")
        _add_source_column(wks, dvdt_sm, f"dV/dt_smooth_trace{trace_num}")

        result_df, ap_count = analyze_trace(time, voltage, dvdt_sm, trace_num, trace_col_1based, settings)
        _write_result_sheet(result_book, trace_num, result_df)

        if ap_count > 0:
            _origin_type(f"Trace {trace_num}: {ap_count} AP(s) detected.")
        else:
            _origin_type(f"Trace {trace_num}: no APs passed the selected filters.")

    _origin_type("Multi-trace AP analysis completed. Results are in Book2.")


if __name__ == "__main__":
    run_ap_analysis(settings_from_labtalk_or_defaults())
```
